# A Geometric Model of Nucleus-Constrained Frustrated Phagocytosis

**DOI:** 10.64898/2026.05.10.724108

**Authors:** Masahiro Fukuda, Jingjiao Guan

## Abstract

Frustrated phagocytosis occurs when phagocytes fail to fully engulf large targets, yet the geometric origins of this physical limit remain poorly defined. Here we present a geometric model that identifies the cell nucleus as an intracellular constraint on engulfment. Extending membrane-limited frameworks, we distinguish an intrinsic phagocytic capacity set by membrane availability from an apparent capacity reduced by nuclear exclusion. Using minimal geometric assumptions, we derive closed-form expressions linking experimentally measurable parameters, including target coverage, volume ratio, and size, to phagocytic capacity and a normalized axial separation that quantifies nuclear accommodation. The model predicts a size- and curvature-independent geometric criterion for nuclear involvement applicable to both spherical and planar targets. These results establish nuclear geometry as a fundamental physical bottleneck in phagocytosis and provide a quantitative framework for interpreting stalled engulfment and nuclear deformation-dependent responses.

## Introduction

Phagocytosis is a fundamental cellular process in which particulate objects typically larger than ∼0.5 µm in diameter are engulfed by specialized cells known as phagocytes and internalized into a membrane-bound compartment called the phagosome.^1,2^ This process is essential for tissue homeostasis, host defense against pathogens, clearance of apoptotic debris, and initiation of adaptive immune responses.^1,2^ Following successful internalization, phagocytosed material is normally degraded within the phagosome through the combined action of hydrolytic enzymes and reactive oxygen species.^3,4^ However, when a target is too large to be fully engulfed, phagocytosis may stall before phagosome closure, resulting in frustrated phagocytosis.^5^ In this state, the phagocytic cup remains open, leading to prolonged cell–target contact and, in some cases, extracellular release of degradative enzymes and reactive species that can damage surrounding tissue.^6,7^ Clinically, frustrated phagocytosis has been implicated in chronic inflammatory diseases such as asbestosis and in adverse reactions to implanted biomaterials.^8–10^ Conversely, the ability of macrophages to engulf whole live tumor cells is emerging as a promising therapeutic strategy in cancer immunotherapy.^11–13^ Given the similar sizes of tumor cells and therapeutic macrophages, the effectiveness of this therapy is likely limited by the number of tumor cells a macrophage can engulf. At the individual cell level, both the occurrence of frustrated phagocytosis and the number of tumor cells that can be engulfed by a macrophage depend on the phagocyte’s phagocytic capacity. Despite its central importance in both pathology and therapy, the physical determinants of phagocytic capacity remain incompletely understood.

Motivated by the hypothesis that phagocytic capacity is governed by membrane availability, we previously developed a general geometric framework to quantify phagocytic capacity during frustrated phagocytosis.^14^ In this model, phagocytic capacity *k* is defined as the ratio of the total membrane area engaged during uptake, including the phagosome and remaining free surface, to the apparent pre-phagocytosis cell surface area, under three physically motivated assumptions: membrane area limitation, surface-minimizing cell geometry, and regulated changes in cellular volume. These assumptions yield closed-form expressions for *k* that depend only on measurable geometric parameters and apply across diverse phagocytic scenarios, including frustrated phagocytosis on flat substrates, engulfment of spherical and disk-shaped objects, and uptake of multiple particles. However, by treating the phagocyte as a mechanically homogeneous body, this framework neglects the presence of mechanically distinct organelles, most notably the nucleus, which is typically the largest and stiffest cellular component.^15^ As a result, the model is most appropriate for phagocytes with highly deformable nuclei, such as neutrophils with multi-lobulated nuclear architecture,^16^ or for uptake of multiple small targets. Recent measurements further show that macrophage membranes can expand more than tenfold locally during phagocytosis, yet engulfment often halts well before this limit is reached, indicating that membrane availability alone cannot explain phagocytic arrest.^17^ Together, these observations suggest that neglecting nuclear geometry and mechanics may lead to an incomplete description of the limits governing frustrated phagocytosis and phagocytic capacity. To address this, we introduce a distinction between an intrinsic phagocytic capacity, defined by membrane availability alone, and an apparent phagocytic capacity inferred from partial engulfment in the presence of geometric constraints such as the nucleus.

During phagocytosis of large objects by macrophages, cells undergo pronounced spreading and global shape remodeling.^18–20^ These deformations necessarily affect the nucleus, which occupies a substantial fraction of the cellular volume and is mechanically stiffer than the surrounding cytoplasm. Analogous nuclear flattening is well documented in non-phagocytic cells that adhere to and spread on stiff planar substrates, where cell spreading imposes geometric confinement on the nucleus.^21^ Such nuclear deformation is a potent regulator of mechanotransduction, most notably through the transcriptional co-activator Yes-associated protein (YAP), whose nuclear localization increases in response to nuclear flattening, cytoskeletal tension, and confinement-induced changes in nuclear envelope geometry.^21^ In macrophages, YAP activation has been linked to inflammatory gene expression and cytoskeletal remodeling, suggesting a mechanistic connection between nuclear deformation and immune function.^22^ These observations raise the possibility that frustrated phagocytosis mechanically couples geometric constraints on engulfment to nuclear deformation–dependent signaling, thereby influencing both phagocytic capacity and downstream cellular responses. Despite this potential significance, existing theoretical descriptions of frustrated phagocytosis do not incorporate the nucleus as a geometric or mechanical constraint. In this context, the present model does not address signaling pathways directly, but instead identifies the geometric conditions under which nuclear deformation–dependent signaling could become mechanically unavoidable.

In addition to membrane availability, the geometry of the phagocyte during frustrated phagocytosis is governed by cortical tension generated by the membrane–cortex composite.^18,23,24^ The actin cortex, mechanically coupled to the plasma membrane, generates an effective surface tension that tends to minimize the area of the free cell surface, such that the free surface of a partially engulfing phagocyte adopts a surface of constant mean curvature, well approximated by a spherical cap.^23^ During the frustrated phagocytosis of a large spherical target or a flat substrate, this area-minimizing tendency reduces the cell height, thereby imposing a compressive load on the nucleus. Given that the nucleus is a viscoelastic rather than rigid body,^25^ and because frustrated phagocytosis typically persists on timescales of minutes to hours,^20^ the pressure generated by cortical tension and geometric confinement can, in principle, significantly deform the nucleus. These mechanical considerations motivate the development of our geometric model of nucleus-constrained frustrated phagocytosis.

## Results

In this model, a phagocyte is assumed to engulf a spherical target and ultimately reach frustrated phagocytosis. Prior to phagocytosis, the cell is taken to be spherical and to contain a spherical nucleus. As phagocytosis proceeds, the cell advances to maximize contact with the target, while the free surface of the cell is assumed to minimize its area. In the absence of a nucleus, this condition leads to a spherical free surface, consistent with our previous geometric model.^14^ Phagocytic progression is constrained by two global factors: a regulated change in cell surface area and a regulated change in cell volume, each characterized by fixed post-to pre-phagocytosis ratios. Engulfment halts when the cell reaches its maximal surface area ratio subject to the prescribed volume ratio. Under these conditions, the geometry of the cell at frustrated phagocytosis is uniquely determined. When a nucleus is present, its finite size and rigidity impose an additional geometric constraint that can limit further engulfment.

Figure 1 illustrates an initially spherical phagocyte containing a spherical nucleus (gray, labeled “Nu.”) interacting with a spherical target, as well as the resulting configuration at frustrated phagocytosis. Upon reaching this state, the target is partially wrapped by the deformed cell. Depending on the size and mechanical rigidity of the nucleus, three distinct geometric cases can arise, as described below.

**Figure 1.**
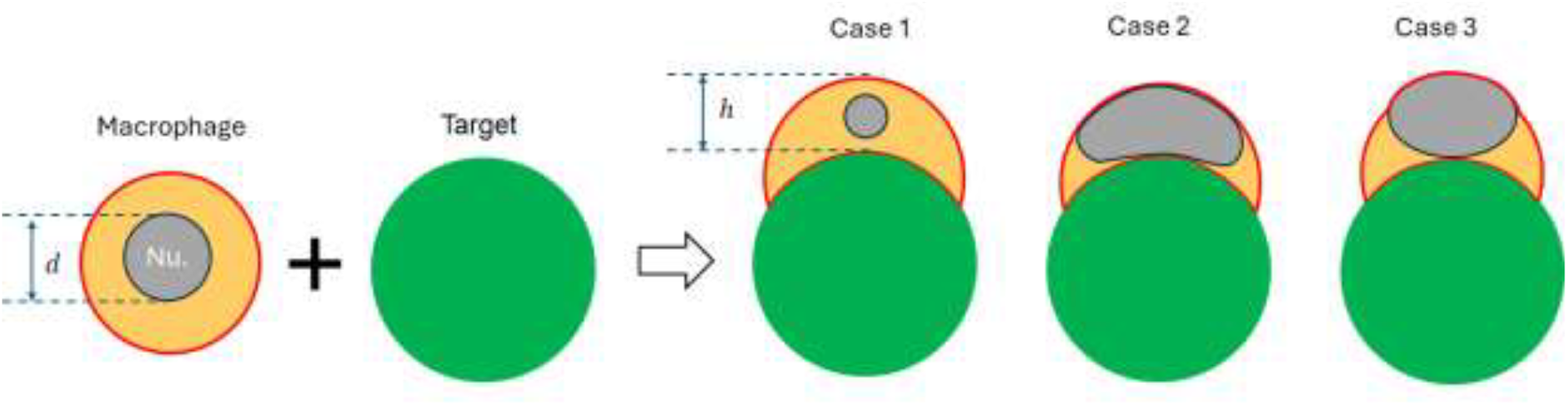
Schematic illustration of nuclear constraints during frustrated phagocytosis of a spherical target by a phagocyte.

In Case 1, the cell has exhausted its available membrane capacity, the free cell surface remains spherical, and the nuclear diameter is smaller than or equal to the distance between the apex of the free cell surface and the apex of the phagocytic target (*d* ≤ *h*). The condition *d* = *h* defines a geometric threshold at which the nucleus simultaneously contacts the apex of the free cell surface and the apex of the phagocytic target. In Case 2, the nucleus is larger than in Case 1 but is assumed to be fully compliant, deforming as needed to accommodate further engulfment. The cell has exhausted its available membrane capacity and the free cell surface remains spherical; this case represents an idealized limit in which the nucleus imposes no constraint on phagocytosis. In Case 3, the nucleus is larger than in Case 1 and mechanically stiffer than the cytoplasm, deforming only upon compression between the advancing free cell surface and the phagocytic target. The cell has exhausted its available membrane capacity, and the portion of the cell surface not in contact with the target deviates from spherical geometry due to nuclear exclusion. As a result, the contact area between the phagocyte and the target is reduced relative to Cases 1 and 2. Case 3 is expected to be the most physiologically relevant scenario for phagocytes with a single globular nucleus, such as macrophages.

For all three cases, the pre-phagocytosis cell is assumed to be spherical, and the post-phagocytosis cell is characterized by a fixed volume ratio relative to the pre-phagocytosis state, although the absolute volumes before and after phagocytosis are not necessarily the same. In Case 1, the nucleus does not deform. In Cases 2 and 3, the nucleus deforms to varying degrees. The phagocyte-target contact area measured in Cases 1 and 2 is expected to be the same and larger than the value measured in Case 3, reflecting the additional geometric constraint imposed by nuclear deformation in the latter case. We therefore denote Cases 1 and 2 as nucleus-unconstrained frustrated phagocytosis and Case 3 as nucleus-constrained frustrated phagocytosis.

Phagocytic capacity, *k*, is traditionally defined as the ratio of the post-phagocytosis cell surface area to the apparent surface area of the spherical pre-phagocytosis cell.^14^ If the post-phagocytosis surface area could be measured directly, *k* could be calculated explicitly for each case. Under the assumptions of the present model, the values of *k* for Cases 1 and 2 are identical and define the intrinsic phagocytic capacity, *k*_*intrinsic*_, whereas the value of *k* for Case 3 is expected to be lower due to nuclear constraint. Because direct measurement of post-phagocytosis surface area for large cell populations is not currently feasible, we instead rely on the experimentally accessible area fraction of the spherical target covered by the phagocyte, σ, which can be readily quantified across large populations, as discussed in the Discussion section.

In this work, we develop a geometric model for frustrated phagocytosis of a spherical target by a phagocyte, in which the free surface of the post-phagocytosis cell is assumed to be spherical. Using this model together with a small set of experimentally measurable parameters, including the target coverage fraction *σ*, we determine an apparent phagocytic capacity, *k*_*apparent*_. Depending on the size and rigidity of the nucleus, *k*_*apparent*_ may be equal to the intrinsic phagocytic capacity, *k*_*intrinsic*_, corresponding to Cases 1 and 2, or smaller than *k*_*intrinsic*_, corresponding to the nucleus-constrained regime (Case 3). If *k*_*intrinsic*_ can be measured independently, comparison between *k*_*apparent*_ and *k*_*intrinsic*_ provides a direct criterion for identifying nuclear constraint, with *k*_*apparent*_ < *k*_*intrinsic*_ indicating nuclear-limited phagocytosis. Thus, the model-derived *k*_*apparent*_ serves as a quantitative metric for assessing the degree of nucleus-constrained frustrated phagocytosis.

In addition, the model enables calculation of the geometric quantity *h* defined in Figure 1 from experimentally measurable parameters, yielding an apparent axial separation, *h*_*apparent*_. This quantity represents the axial space demanded by the engulfment geometry at the stalled phagocytic state. Comparison of *h*_*apparent*_ with the diameter of the initially spherical nucleus provides a geometric criterion for nuclear involvement: when *h*_*apparent*_ < *d*, continued engulfment would require either nuclear deformation or arrest of membrane advancement, depending on nuclear rigidity. Thus, *h*_*apparent*_ characterizes geometric incompatibility imposed by frustrated phagocytosis rather than directly measuring nuclear deformation. Importantly, *h*_*apparent*_ is a geometric demand imposed by engulfment rather than a direct measure of nuclear deformation; a perfectly rigid nucleus may satisfy *h*_*apparent*_ < *d* without deforming, whereas a compliant nucleus may deform even when this inequality is only marginally violated.

Finally, the model enables prediction of whether the engulfment geometry is compatible with accommodating the nucleus for a phagocyte with a known intrinsic phagocytic capacity, *k*_*intrinsic*_, on either convex spherical or flat targets.

Geometries of the model are shown in Figure 2. A phagocyte with effective radius *r* (yellow) partially engulfs a spherical target of radius *x* (white). During frustrated phagocytosis, the free surface of the phagocyte adopts a spherical-cap geometry that wraps around the target. In the composite geometry (right), *R* denotes the radius of curvature of the phagocyte free surface, *x* the target radius, and *a* represents the radial distance from the symmetry axis to the circle of intersection between the spherical target and the spherical cap. The quantities *h*_1_ and *h*_2_ denote the axial distances from the plane of intersection between the target and the phagocyte surface to the apex of the spherical target and to the apex of the phagocyte free surface, respectively. Dashed lines indicate geometric reference axes and distances used in the derivation of the governing equations.

**Figure 2.**
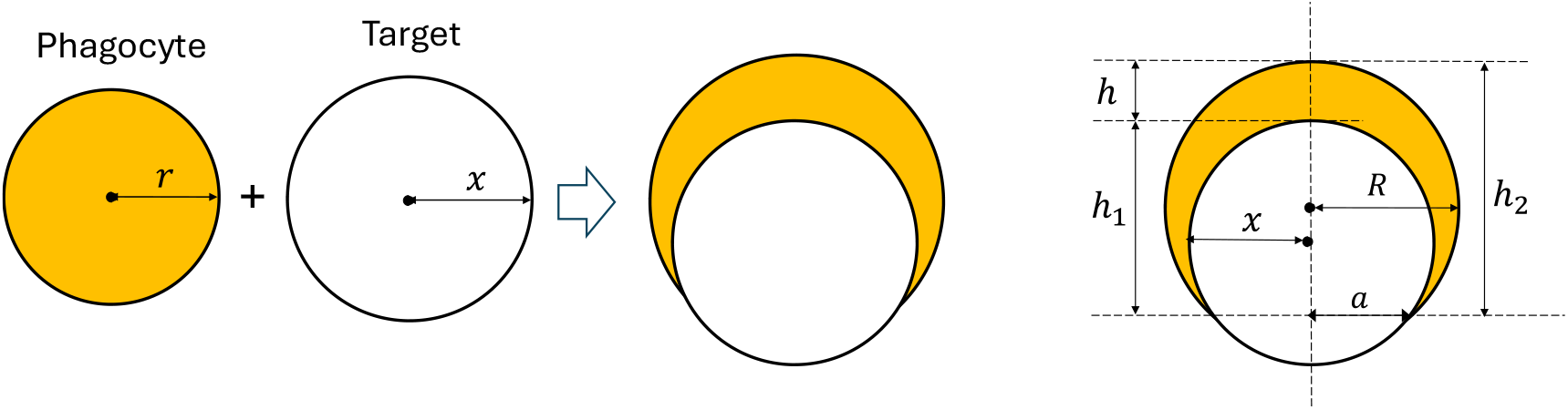
Geometric definitions for frustrated phagocytosis of a spherical target by a phagocyte.

Based on the geometry shown in Figure 2, we derive the following governing relations using the Pythagorean theorem:

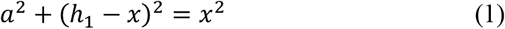

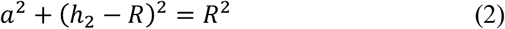

We define the fraction of the spherical target surface covered by the cell as *σ*, which can be experimentally measured, for example, by flow cytometry:

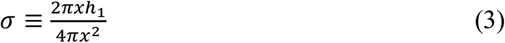

Following our previous work,^14^ we define the volume ratio between the post-phagocytosis cell and the pre-phagocytosis cell as *ε*:

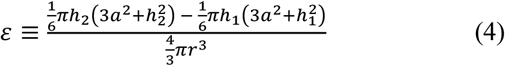

We further define the surface area ratio between the post-phagocytosis cell and the pre-phagocytosis cell as *k*, consistent with our earlier formulation: ^14^

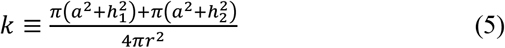

Finally, we define the vertical distance between the apex of the spherical target and the apex of the cell as

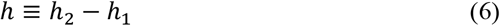

which quantifies the available axial space within the cell for accommodating the nucleus.

Based on Eqs. (1)–(6), our primary objective is to derive explicit closed-form expressions for biologically meaningful parameters in terms of experimentally measurable variables. The biologically meaningful parameters include the phagocytic capacity *k*, and the normalized axial separation *h*/*r*, which quantifies the space available to accommodate the nucleus. The experimentally measurable variables are the normalized target radius *x*/*r*, the target coverage fraction *σ*, and the volume ratio *ε*. A secondary objective is to identify hidden geometric relationships among *h*/*r, k*, and *ε*.

To derive explicit expressions for *k* and *h*/*r*, we first obtain a closed-form solution for *h*_2_/*r*. Using Eqs. (1), (3), and (4), we arrive at the following cubic equation:

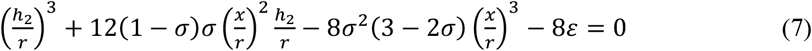

The discriminant of this cubic equation is

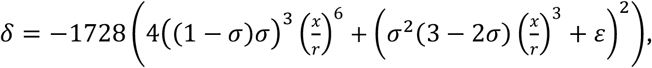

which is strictly negative for 0 ≤ *σ* ≤ 1. Consequently, Eq. (7) admits a single real solution and a pair of complex conjugate solutions, indicating that for a given target size, coverage fraction, and volume ratio, the geometry of frustrated phagocytosis is uniquely determined. The real root is given by

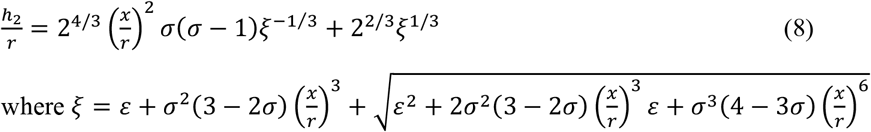

By combining Eq. (8) with Eqs. (1), (3) and (5), we derive a closed-form expression for the phagocytic capacity *k* (Eq. (9)). When the target radius *x*, cell radius *r*, and coverage fraction *σ* are experimentally measured, this expression can be used to obtain the apparent phagocytic capacity *k*_*apparent*_.

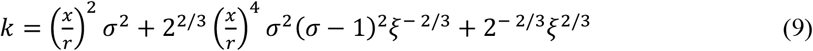

where *ξ* is defined in (8).

Equation (9) is plotted in Figure 3 as a function of the normalized target radius *x*/*r* and the target coverage fraction *σ*. Figure 3A shows the resulting three-dimensional surface, illustrating how geometric engagement of a spherical target and the extent of coverage jointly determine membrane utilization during frustrated phagocytosis. Figure 3B presents a one-dimensional slice of this surface at a representative target size (*x*/*r* = 2), demonstrating the monotonic increase of the apparent phagocytic capacity *k* with increasing coverage fraction. Notably, although the closed-form expression for *k* is nonlinear, the physically relevant branch exhibits an approximately linear dependence on target coverage over the experimentally relevant range, reflecting the uniform geometric contribution of incremental coverage to membrane utilization.

**Figure 3.**
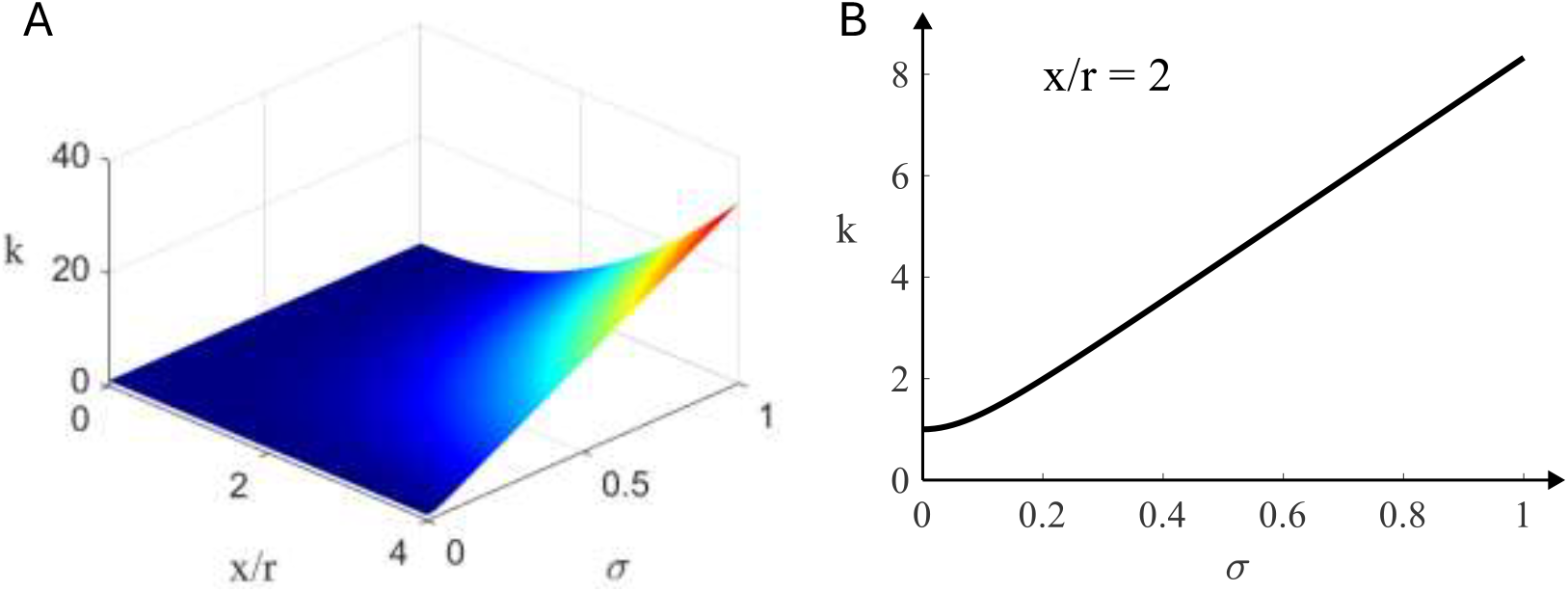
Dependence of phagocytic capacity on target size and coverage. (A) Three-dimensional surface plot of the apparent phagocytic capacity *k* as a function of the normalized target radius *x*/*r* and the target coverage fraction *σ*, as given by Eq. (9). (B) One-dimensional slice of the surface at a representative target size (*x*/*r* = 2), illustrating the monotonic increase of *k* with increasing coverage fraction *σ*.

By combining Eq. (8) with Eqs. (3) and (6), we obtain the following closed-form expression for the normalized axial separation *h*/*r*:

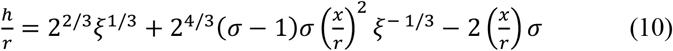

where *ξ* is defined in Eq. (8).

Equation (10) is visualized in Figure 4 as a function of the normalized target radius *x*/*r* and the target coverage fraction *σ*. The resulting surface plot shows that, for a fixed target size *x*/*r*, the normalized axial separation *h*/*r* decreases monotonically with increasing target coverage. This behavior is further illustrated in the one-dimensional slice shown in panel B, which highlights the monotonic reduction of *h*/*r* with increasing *σ* at a representative target size. Together, these plots demonstrate that increased target coverage drives a progressive reduction in the axial separation required by the engulfment geometry. Accordingly, in a frustrated phagocytosis experiment, measurement of *x, r, σ* and *ε* enables calculation of the apparent axial separation *h*_*apparent*_ using Eq. (10).

**Figure 4.**
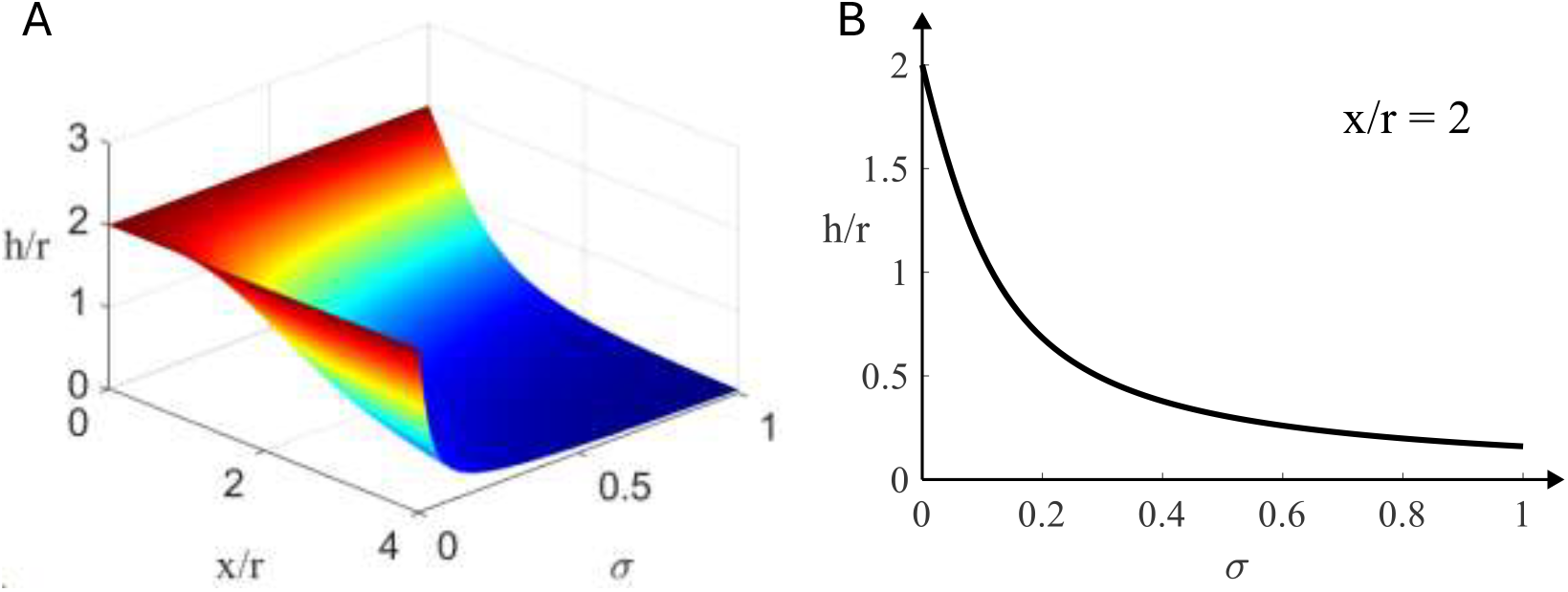
Dependence of the normalized axial separation *h*/*r* on target size and coverage. (A) Three-dimensional surface plot of Eq. (10) showing *h*/*r* as a function of the normalized target radius *x*/*r* and the target coverage fraction *σ*. For a fixed target size, increasing coverage leads to a monotonic decrease in *h*/*r*. (B) One-dimensional slice of the surface at a representative target size (*x*/*r* = 2), illustrating the monotonic dependence of *h*/*r* on coverage fraction *σ*.

Our model enables the establishment of a theoretical threshold for nuclear deformation in a phagocyte with a given *k*_*intrinsic*_ and *ε*. Based on Cases 1 and 2 in Figure 1, the nucleus does not deform when *d* ≤ *h*, which is equivalently expressed as *d*/*r* ≤ *h*/*r*. Because both *d* and *r* are experimentally measurable quantities, it is therefore useful to derive an equation that relates the dimensionless parameter *h*/*r* to the membrane capacity *k* and the geometric parameter *ε*.

By combining Eqs. (4) and (5), we eliminate the intermediate variables *h*_1_, *h*_2_, and *a*, yielding the following cubic equation in terms of *h*/*r, k*, and *ε*:

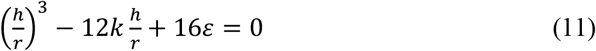

The discriminant of this equation is *δ* = 6912(*k*^3^ − *ε*^2^). The condition *ε*^2^ = *k*^3^ corresponds to the spherical limit, whereas *ε*^2^ < *k*^3^ spans all non-spherical shapes. The inequality *ε*^2^ ≤ *k*^3^ therefore defines the set of physically admissible geometries. When *δ* = 0 (except for the trivial case *ε* = *k* = 0), Eq. (11) admits three real solutions, two of which are equal. When *δ* > 0, there are three distinct real solutions, a regime that is directly relevant to frustrated phagocytosis in phagocytes. For mathematical completeness, when *δ* < 0, the equation yields one real solution and a pair of complex conjugate solutions.

Equation (11) is plotted in Figure 5A as a function of the phagocytic capacity *k* and the volume ratio *ε*. The parameter *k* is varied from 0 to 6 and *ε* from 0 to 2, ranges chosen to encompass all physically relevant conditions. Within this domain, Eq. (11) admits three real solutions, one of which is always negative and therefore physically inadmissible. The remaining two solutions are strictly positive and define two distinct geometric branches. The red curve, indicated by an arrow, marks the locus at which these two positive solutions converge. This curve is defined by *ε*^2^ = *k*^3^ and 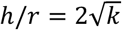, and geometrically corresponds to the limiting case in which the cell remains spherical at the stalled state of frustrated phagocytosis.

**Figure 5.**
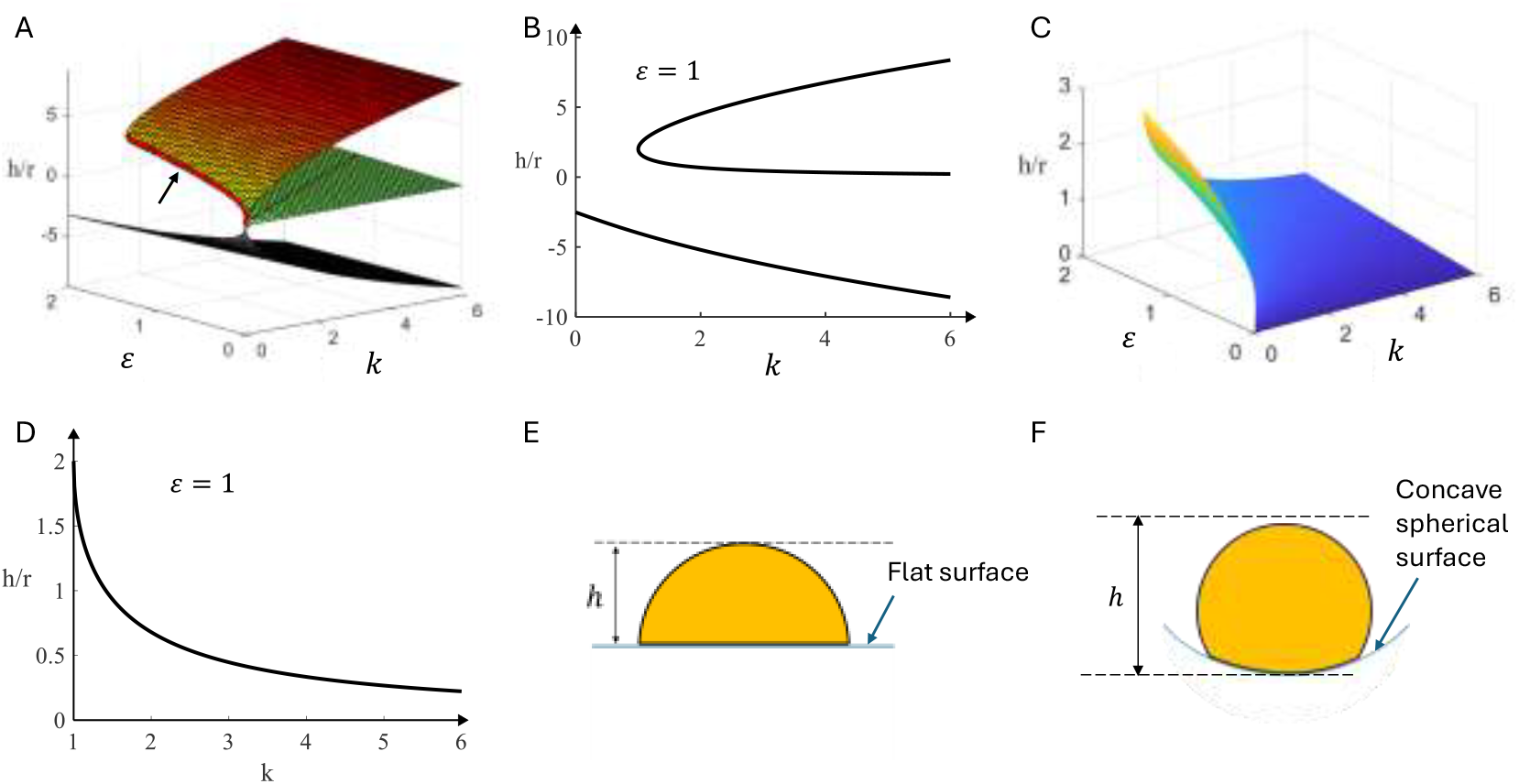
Geometric dependence of the normalized axial separation *h*/*r* on phagocytic capacity *k* and volume ratio *ε*, and the resulting criterion for nuclear involvement. (A) Three-dimensional solution surface showing *h*/*r* as a function of *k* and *ε*. (B) Planar slice of the solution surface at *ε* = 1, revealing two positive solution branches: a physically relevant branch in which *h*/*r* decreases monotonically with increasing *k*, and an unphysical branch in which *h*/*r* increases with *k*. (C) Three-dimensional rendering of the admissible solution surface generated using Eq. (12). (D) Dependence of the normalized axial separation *h*/*r* on *k* at *ε* = 1. (E) Schematic of frustrated phagocytosis on a flat surface, defining the axial separation *h*. (F) Corresponding schematic for a concave spherical surface.

For any fixed value of *ε*, the two positive solutions exhibit opposite monotonic trends with increasing *k*: the upper surface increases with *k*, whereas the lower surface decreases with *k*. This structure is further clarified in Figure 5B, which shows a planar slice (top view) of the three-dimensional solution surface at *ε* = 1, corresponding to the case in which the cell maintains constant volume before and after phagocytosis. The slice reveals two positive solution branches with opposite monotonic behavior: one branch decreases monotonically with increasing *k*, while the other increases with *k*. Because an increase in membrane capacity is expected to permit further engulfment before stalling, the physically relevant solution for phagocytosis is the branch for which the normalized axial separation *h*/*r* decreases with increasing *k*. In contrast, the branch in which *h*/*r* increases with *k* corresponds to a geometric configuration that is not accessed during phagocytosis and is therefore excluded from further consideration.

Accordingly, the phagocytosis-relevant solution is given by

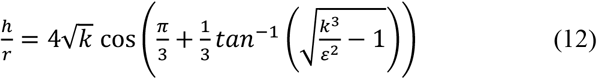

Equation (12) is plotted in Figure 5C as a function of *k* and *ε*. For clarity, Figure 5D shows *h*/*r* as a function of *k* at *ε* = 1, illustrating that *h*/*r* decreases monotonically with increasing *k*.

If a macrophage’s intrinsic phagocytic capacity *k*_*intrinsic*_ and volume ratio *ε* are known, Eq. (12) enables direct calculation of the corresponding normalized axial separation *h*/*r*, which can be compared with the normalized nuclear diameter *d*/*r*. When *h*/*r* > *d*/*r*, the engulfment geometry does not impose axial confinement on the nucleus, and the free cell surface can remain spherical without requiring nuclear deformation. Conversely, when *h*/*r* < *d*/*r*, the advancing membrane imposes a geometric compression on the nucleus. Because the nucleus is a viscoelastic body and frustrated phagocytosis typically persists over extended timescales,^20,25^ this geometric incompatibility is expected to result in time-dependent nuclear deformation, the extent of which depends on both the mechanical properties of the nucleus and the stresses generated by cortical tension.

To our knowledge, no experiment has been specifically designed to measure *k*_*intrinsic*_ while experimentally excluding or minimizing nuclear constraints. As a result, reliable values of *k*_*intrinsic*_ have not been reported for most macrophage types. However, based on published data for two-dimensional frustrated phagocytosis and our previous geometric model, we determined an apparent phagocytic capacity *k*_*apparent*_ = 5.17 for murine bone marrow–derived macrophages (BMDMs).^14^ Using this value and assuming *ε* = 1, Eq. (12) yields *h*/*r* ≈ 0.26.

Swanson *et al*. reported an aqueous nucleocytoplasmic volume ratio of approximately 0.18 in murine BMDMs,^26^ corresponding to a nuclear volume fraction of approximately 0.15 and, under a spherical approximation, to a nucleus-to-cell diameter ratio of *d*/*r* ≈ 1.06. Because *k*_*apparent*_ is expected to be lower than *k*_*intrinsic*_, and because *h*/*r* decreases monotonically with increasing *k*, the values of *h*/*r* calculated using *k*_*intrinsic*_ must be even smaller than those obtained from *k*_*apparent*_. Consequently, because *h*/*r* ≈ 0.26 is substantially smaller than *d*/*r* ≈ 1.06, nuclear deformation is expected to occur during phagocytosis in the macrophage type considered here.

Notably, Eq. (11) does not contain the target radius *x*, indicating that *h*/*r* depends only on *k* and *ε* and is independent of the size of the phagocytic object. Moreover, even when the object is a flat surface and *h* is defined as shown in Figure 5E, the same form of Eq. (11) can be derived. Because flat substrates are commonly used as the phagocytic target in studies of frustrated phagocytosis known as two-dimensional (2D) frustrated phagocytosis, this result demonstrates the feasibility of predicting nuclear deformation in both spherical- and planar-target geometries when *k* and *ε* are known.

Moreover, the independence of Eq. (11) from the target radius indicates that whether nuclear deformation occurs is insensitive to the size of the spherical phagocytic target and to whether the target is spherical or planar. In other words, the onset of nuclear deformation is independent of the curvature of the phagocytic object. Finally, even when the substrate is a concave spherical surface and *h* is defined as shown in Figure 5F, the same form of Eq. (11) can be derived, demonstrating that this geometric criterion is invariant to the sign (polarity) of substrate curvature.

Step-by-step derivations of Eqs. (1)-(12), including generalizations of Eq. (11) to flat and concave spherical targets, are provided in the Supplementary Information.

## Discussion

This work identifies the cell nucleus as a fundamental geometric constraint that limits frustrated phagocytosis independently of biochemical regulation. By explicitly incorporating nuclear exclusion into a geometric framework, we show that partial engulfment can arise purely from spatial incompatibility between the advancing phagocytic cup and the intracellular nucleus, even when membrane area and volume would otherwise permit further engulfment. The model defines intrinsic and nucleus-constrained phagocytic capacity as distinct regimes and introduces the normalized axial separation as a measurable geometric metric of nuclear involvement. Importantly, the predicted onset of nuclear deformation depends on intrinsic cellular geometry rather than target size or curvature, implying that the limits of phagocytosis are set primarily by intracellular organization. Together, these results establish nuclear geometry as a previously underappreciated determinant of phagocytic capacity and provide a quantitative basis for linking frustrated phagocytosis to nuclear deformation–dependent cellular responses. More broadly, this framework illustrates how intracellular geometry can impose physical limits on cellular functions that are often interpreted primarily through molecular regulation.

From a methodological perspective, this study illustrates how a complex biological process can be systematically reduced to an exact geometric and algebraic formulation. Frustrated phagocytosis is first abstracted as a minimal geometric problem by representing the phagocyte, target, and nucleus with spherical geometries subject to conservation of membrane area and cell volume. These biologically motivated constraints generate a closed set of nonlinear algebraic relations, which reduce to cubic equations that encode the compatibility between external engulfment geometry and internal nuclear exclusion. Exact closed-form solutions of these cubic equations yield explicit expressions for biologically meaningful quantities, including phagocytic capacity and axial nuclear clearance, directly in terms of experimentally measurable parameters. This progression from biology to geometry, from geometry to algebra, and back to biological interpretation demonstrates that the limits of phagocytosis can be governed by precise mathematical structure, revealing how intracellular organization imposes fundamental geometric bounds on cellular function.

To translate this geometric framework into an experimentally testable reference point, it is necessary to determine the intrinsic phagocytic capacity under conditions in which nuclear constraints are effectively eliminated. We therefore propose measuring macrophage uptake of multiple small spherical targets whose individual sizes are insufficient to induce nuclear deformation, as schematically illustrated in Figure 6. By feeding macrophages with a large number of uniformly small (e.g., 1-µm-diameter), IgG-opsonized microspheres, membrane area engaged in phagocytosis can be driven to saturation while preserving an approximately spherical cell morphology and minimizing nuclear involvement, such that uptake is limited solely by membrane availability. In practice, fluorescent microspheres will be incubated with macrophages and analyzed by flow cytometry, with trypan blue used to quench fluorescence from extracellularly bound particles. Because the microspheres are monodisperse, fluorescence intensity per cell scales linearly with the number of internalized particles, enabling determination of the number of phagocytosed microspheres per cell, *n*. The intrinsic phagocytic capacity, *k*_*intrinsic*_, can then be calculated directly using Eq. (13), as derived previously^14^ and detailed in the Supplementary Information, and used as a reference against which *k*_*apparent*_ measured during frustrated phagocytosis of large targets can be compared to quantify the extent of nuclear constraint.

**Figure 6.**
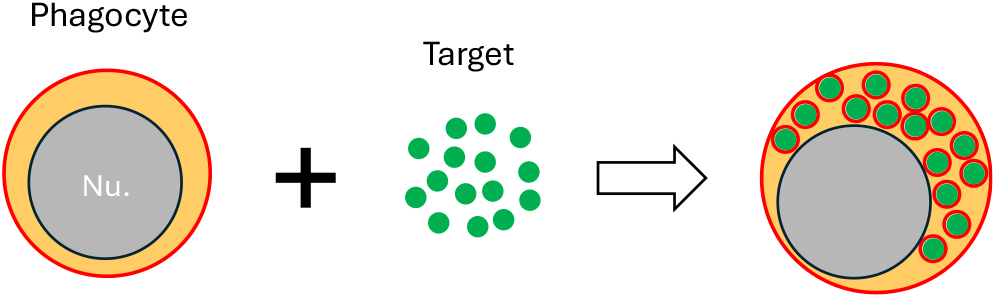
Proposed method for determining *k*_*intrinsic*_ through phagocytosis of multiple small spherical targets by a phagocyte.

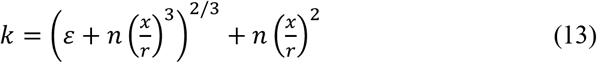

Having established an experimental strategy for determining the intrinsic phagocytic capacity, the same geometric framework can be applied to quantify nucleus-constrained frustrated phagocytosis. To do so, the required parameters can be obtained by combining flow cytometry and confocal microscopy. Flow cytometry can be used to quantify the target coverage fraction by labeling exposed target surfaces after frustrated phagocytosis, enabling population-level measurement of partial engulfment, as demonstrated in classic studies of macrophage phagocytic capacity.^20^ In parallel, confocal microscopy of suspended and frustrated macrophages allows three-dimensional reconstruction of cell volumes before and after phagocytosis, from which the volume ratio can be extracted.^27,28^ Together with independently measured target size and cell radius, these experimentally accessible quantities provide all necessary inputs to compute phagocytic capacity and axial nuclear clearance using the closed-form expressions derived here.

Several limitations of the present framework should be noted. The model is intentionally static and geometric and therefore does not capture the dynamic processes underlying phagocytosis, including time-dependent membrane trafficking, cytoskeletal remodeling, or the viscoelastic response of the nucleus. In addition, the assumption of spherical symmetry for the phagocyte, nucleus, and target simplifies analysis but does not account for nuclear eccentricity, off-center positioning, or anisotropic cytoskeletal organization observed in vivo. These simplifications are deliberate, as the goal of the model is to identify geometric constraints that apply independent of molecular detail. Deviations from spherical geometry or homogeneous mechanics may shift quantitative predictions but are unlikely to alter the qualitative conclusion that nuclear exclusion imposes a fundamental geometric limit on phagocytic capacity.

This work lays a foundation for extending the present geometric framework through coordinated experimental and theoretical efforts. Experimentally, time-resolved microscopy could be used to quantify the evolving shapes and volumes of both the whole cell and its nucleus during stalled engulfment, providing direct tests of the geometric constraints identified here. On the theoretical side, further development of the model may reveal underappreciated connections between geometric features and biologically meaningful cellular properties, including membrane tension, cytosolic osmotic pressure, and deformation of the cytoskeleton and endoplasmic reticulum. Together, these directions position geometry as a unifying lens for understanding frustrated phagocytosis.

Although developed here in the context of frustrated phagocytosis, the geometric constraint identified by this model reflects a more general principle that applies whenever cells undergo large-scale shape remodeling under surface-area and volume constraints in the presence of a relatively large nucleus. Under such conditions, global changes in cell geometry impose spatial demands that can force the nucleus to deform or otherwise limit further advancement of the cell boundary. Similar geometric incompatibilities may therefore arise during processes such as confined migration through narrow constrictions during monocyte extravasation, wrapping of axons by Schwann cells, or partial wrapping of blood vessels by pericytes, where axial or spatial clearance of the nucleus becomes a limiting factor. In this sense, the present framework provides a geometric lens for distinguishing cellular behaviors limited by intrinsic spatial incompatibility from those limited by molecular regulation, highlighting nuclear geometry as a fundamental determinant of cellular outcomes during extreme shape change.

## Conclusions

We present a geometric model of frustrated phagocytosis that explicitly accounts for constraints imposed by the cell nucleus. By analyzing limiting cases of a nucleus that is either highly deformable or effectively rigid, we distinguish an intrinsic phagocytic capacity, set by membrane availability alone, from a nucleus-constrained apparent phagocytic capacity that reflects geometric exclusion by the nucleus, and we show how nuclear geometry reduces the apparent capacity relative to its intrinsic limit. The model introduces a simple geometric metric that provides a direct, experimentally accessible criterion for nuclear involvement during frustrated phagocytosis. Together, these results identify the nucleus as a fundamental geometric bottleneck in phagocytosis and provide a minimal framework for linking nuclear geometry and mechanics to limits on cellular engulfment.

