## Supplementary Information for "A Geometric Model of Nucleus-Constrained Frustrated Phagocytosis"

**Table of Contents:**

Overview

Equations (1) and (2): Pythagorean Geometric Relations

Equation (3): Target Coverage Fraction σ

Equation (4): Volume Ratio ε

Equation (5): Surface Area Ratio (Phagocytic Capacity) k

Equation (6): Axial Separation h

Equation (7): Cubic Equation for $h_2/r$

Equation (8): Closed-Form Solution for $h_2/r$

Equation (9): Phagocytic Capacity k

Equation (10): Normalized Axial Separation h/r

Equation (11): Cubic Relation among h/r, k, and ε

Generalization of Eq. (11) to a Flat Target

Generalization of Eq. (11) to a Concave Spherical Target

Equation (12): Physically Relevant Root of Eq. (11)

Equation (13): Phagocytic Capacity for Multiple Small Targets

Summary Table of Derived Equations

### Overview

This document provides step-by-step derivations of all governing equations (Eqs. 1–13) presented in the manuscript. The model geometry is defined by two intersecting spherical caps: a phagosome cap of radius $x$ (the target) and a free-surface cap of radius $R$ (the phagocyte), sharing a common circular intersection of radius $a$. The axial heights from the intersection plane to the target apex and free-surface apex are $h_1$ and $h_2$, respectively. The pre-phagocytosis cell is a sphere of radius $r$.

**Summary of notation:**

- $r$: effective radius of the pre-phagocytosis (spherical) cell
- $x$: radius of the spherical target
- $R$: radius of curvature of the free cell surface (spherical cap)
- $a$: radius of the contact circle at the intersection plane
- $h_1$: axial distance from intersection plane to target apex
- $h_2$: axial distance from intersection plane to apex of free cell surface
- $\sigma$: fraction of target surface covered by the phagocyte
- $\varepsilon$: ratio of post- to pre-phagocytosis cell volume
- $k$: ratio of post- to pre-phagocytosis cell surface area (phagocytic capacity)
- $h = h_2 - h_1$: axial separation available for nuclear accommodation (for the convex spherical target; redefined as $h = h_1 + h_2$ for the concave spherical target generalization)

**Equations (1) and (2): Pythagorean Geometric Relations**

The intersection circle of radius $a$ lies in a common plane. Each of the two spherical caps meets this circle at its base.

**Equation (1): Target sphere**

The target is a sphere of radius $x$. Its center lies at distance $x$ from the sphere's own apex. The axial distance from the intersection plane to the target apex is $h_1$, so the center of the target sphere lies at axial distance $(h_1 - x)$ from the intersection plane (measured in the direction of the target apex). Applying the Pythagorean theorem to the right triangle formed by the contact radius $a$, the axial offset of the center $(h_1 - x)$, and the target radius $x$:

$$a^2 + (h_1 - x)^2 = x^2.$$

Note that $(h_1 - x)^2 = (x - h_1)^2$, so this is symmetric in sign.

**Equation (2): Free cell surface**

The free surface of the phagocyte is a spherical cap of radius $R$. Its center lies at axial distance $(h_2 - R)$ from the intersection plane. Applying the Pythagorean theorem:

$$a^2 + (h_2 - R)^2 = R^2.$$

**Equation (3): Target Coverage Fraction $\sigma$**

The lateral surface area of a spherical cap of height $h_1$ on a sphere of radius $x$ is $2\pi x h_1$ (standard formula for spherical cap area). The total surface area of the target sphere is $4\pi x^2$. Therefore, the fraction covered is:

$$\sigma \equiv \frac{2\pi x h_1}{4\pi x^2} = \frac{h_1}{2x}.$$

Equivalently, $h_1 = 2\sigma x$, a relation used throughout subsequent derivations.

**Equation (4): Volume Ratio $\varepsilon$**

The post-phagocytosis cell volume is bounded by the free-surface cap above and the phagosome cap below, both sharing the intersection circle of radius $a$.

**Volume of a spherical cap**

For a spherical cap of height $h$ and base radius $a$, the volume is:

$$V_{\text{cap}} = \frac{\pi h(3a^2 + h^2)}{6}.$$

(This is equivalent to $\pi h^2(3R - h)/3$ via the identity $a^2 = h(2R - h)$, where $R$ is the sphere radius.)

**Cell volume**

The post-phagocytosis cell occupies the region bounded by: (i) the spherical cap of the free surface (radius $R$, height $h_2$, from the intersection plane to the apex of the free surface), and (ii) the spherical cap of the target (radius $x$, height $h_1$, from the intersection plane to the target apex, representing the phagosome).

The net cell volume is obtained by subtracting the inner phagosome cap from the outer free-surface cap:

$$V_{\text{cell}} = \frac{\pi h_2(3a^2 + h_2^2)}{6} - \frac{\pi h_1(3a^2 + h_1^2)}{6}.$$

The pre-phagocytosis cell is a sphere of radius $r$ with volume $4/3\,\pi r^3$. Therefore:

$$\varepsilon \equiv \frac{\frac{1}{6}\pi h_2(3a^2 + h_2^2) - \frac{1}{6}\pi h_1(3a^2 + h_1^2)}{\frac{4}{3}\pi r^3}.$$

**Equation (5): Surface Area Ratio (Phagocytic Capacity) $\boldsymbol{k}$**

**Surface area of a spherical cap**

The curved surface area of a spherical cap of height $h$ and base radius $a$ on a sphere of radius $R$ is $2\pi Rh$. Using $a^2 = h(2R - h)$, this simplifies to:

$$S_{\text{cap}} = 2\pi Rh = \pi(a^2 + h^2),$$

since $a^2 + h^2 = h(2R - h) + h^2 = 2Rh.$

**Post-phagocytosis surface**

The total post-phagocytosis cell surface consists of:

- Phagosome (cap of target sphere, height $h_1$): $\pi(a^2 + h_1^2)$
- Free cell surface (cap of free sphere, height $h_2$): $\pi(a^2 + h_2^2)$

The pre-phagocytosis surface area is $4\pi r^2$. Therefore:

$$k \equiv \frac{\pi(a^2 + h_1^2) + \pi(a^2 + h_2^2)}{4\pi r^2}.$$

**Equation (6): Axial Separation $h$**

The quantity $h$ is defined as the axial distance between the apex of the target and the apex of the free cell surface:

$$h \equiv h_2 - h_1.$$

This represents the internal axial space available to accommodate the nucleus at the stalled phagocytic state.

**Equation (7): Cubic Equation for $h_2/r$**

**Step 1: Simplify $a^2$ and $h_1$ using $\sigma$**

From Eq. (1): $a^2 = x^2 - (h_1 - x)^2 = 2xh_1 - h_1^2$.
From Eq. (3): $h_1 = 2\sigma x$. Substituting:

$$a^2 = 2x(2\sigma x) - (2\sigma x)^2 = 4\sigma x^2 - 4\sigma^2 x^2 = 4\sigma(1-\sigma)x^2.$$

**Step 2: Compute $h_1(3a^2 + h_1^2)$**

$$\begin{aligned} h_1(3a^2 + h_1^2) &= 2\sigma x(3 \cdot 4\sigma(1-\sigma)x^2 + (2\sigma x)^2) \\ &= 2\sigma x \cdot x^2(12\sigma(1-\sigma) + 4\sigma^2) \\ &= 2\sigma x^3(12\sigma - 12\sigma^2 + 4\sigma^2) \\ &= 2\sigma x^3 \cdot (12\sigma - 8\sigma^2) \\ &= 8\sigma^2(3-2\sigma)x^3. \end{aligned}$$

**Step 3: Substitute into Eq. (4)**

Let $V = h_2(3a^2 + h_2^2)$. Then:

$$\begin{aligned} 8r^3\varepsilon &= h_2(3a^2 + h_2^2) - 8\sigma^2(3-2\sigma)x^3 \\ &= h_2(12\sigma(1-\sigma)x^2 + h_2^2) - 8\sigma^2(3-2\sigma)x^3 \\ &= 12\sigma(1-\sigma)x^2 h_2 + h_2^3 - 8\sigma^2(3-2\sigma)x^3. \end{aligned}$$

**Step 4: Non-dimensionalize**

Dividing through by $r^3$:

$$8\varepsilon = 12\sigma(1-\sigma)\left(\frac{x}{r}\right)^2 \frac{h_2}{r} + \left(\frac{h_2}{r}\right)^3 - 8\sigma^2(3-2\sigma)\left(\frac{x}{r}\right)^3.$$

Rearranging:

$$\left(\frac{h_2}{r}\right)^3 + 12(1-\sigma)\sigma\left(\frac{x}{r}\right)^2 \frac{h_2}{r} - 8\sigma^2(3-2\sigma)\left(\frac{x}{r}\right)^3 - 8\varepsilon = 0.$$

This is a *depressed cubic* in $h_2/r$ of the form $t^3 + pt + q = 0$ with:

$$p = 12(1-\sigma)\sigma\left(\frac{x}{r}\right)^2, \qquad q = -8\sigma^2(3-2\sigma)\left(\frac{x}{r}\right)^3 - 8\varepsilon.$$

**Discriminant**

The discriminant of $t^3 + pt + q = 0$ is $\delta = -4p^3 - 27q^2$. Substituting:

$$
\begin{aligned}
\delta \quad &= -4[12(1-\sigma)\sigma]^3\left(\frac{x}{r}\right)^6 - 27[8\sigma^2(3-2\sigma)\left(\frac{x}{r}\right)^3 + 8\varepsilon]^2 \\
&= -6912(1-\sigma)^3\sigma^3\left(\frac{x}{r}\right)^6 - 1728[\sigma^2(3-2\sigma)\left(\frac{x}{r}\right)^3 + \varepsilon]^2 \\
&= -1728\left[4((1-\sigma)\sigma)^3\left(\frac{x}{r}\right)^6 + \left(\sigma^2(3-2\sigma)\left(\frac{x}{r}\right)^3 + \varepsilon\right)^2\right].
\end{aligned}
$$

Since both terms inside the bracket are non-negative for $0 \leq \sigma \leq 1$, we have $\delta < 0$, guaranteeing exactly one real root.

**Equation (8): Closed-Form Solution for $h_2/r$**

**Cardano's formula**

For the depressed cubic $t^3 + pt + q = 0$ with one real root, Cardano's formula gives:

$$t = u + v, \quad u = \sqrt[3]{-\frac{q}{2} + \sqrt{\frac{q^2}{4} + \frac{p^3}{27}}}, \quad v = \sqrt[3]{-\frac{q}{2} - \sqrt{\frac{q^2}{4} + \frac{p^3}{27}}},$$

with $uv = -p/3$.

**Compute $-q/2$ and $q^2/4 + p^3/27$**

$$-\frac{q}{2} = 4\sigma^2(3 - 2\sigma)\left(\frac{x}{r}\right)^3 + 4\varepsilon = 4\left[\sigma^2(3 - 2\sigma)\left(\frac{x}{r}\right)^3 + \varepsilon\right].$$

$$\frac{q^2}{4} + \frac{p^3}{27} = 16\left[\sigma^2(3 - 2\sigma)\left(\frac{x}{r}\right)^3 + \varepsilon\right]^2 + 64\,\sigma^3(1 - \sigma)^3\left(\frac{x}{r}\right)^6.$$

Taking the square root:

$$\sqrt{\frac{q^2}{4} + \frac{p^3}{27}} = 4\sqrt{\left[\sigma^2(3 - 2\sigma)\left(\frac{x}{r}\right)^3 + \varepsilon\right]^2 + 4\,\sigma^3(1 - \sigma)^3\left(\frac{x}{r}\right)^6}.$$

**Define $\xi$**

$$\begin{aligned} u^3 &= -\frac{q}{2} + \sqrt{\frac{q^2}{4} + \frac{p^3}{27}} \\ &= 4\left[\varepsilon + \sigma^2(3 - 2\sigma)\left(\frac{x}{r}\right)^3 + \sqrt{\varepsilon^2 + 2\sigma^2(3 - 2\sigma)\left(\frac{x}{r}\right)^3\varepsilon + \sigma^3(4 - 3\sigma)\left(\frac{x}{r}\right)^6}\right] = 4\xi, \end{aligned}$$

where the expression inside the outer square root was simplified by expanding the left-hand side and collecting the coefficient of $(x/r)^6$:

$$\begin{aligned} &\left[\sigma^2(3 - 2\sigma)\left(\frac{x}{r}\right)^3 + \varepsilon\right]^2 + 4\sigma^3(1 - \sigma)^3\left(\frac{x}{r}\right)^6 \\ &\quad = \varepsilon^2 + 2\sigma^2(3 - 2\sigma)\left(\frac{x}{r}\right)^3\varepsilon + [\sigma^4(3 - 2\sigma)^2 + 4\sigma^3(1 - \sigma)^3]\left(\frac{x}{r}\right)^6. \end{aligned}$$

The coefficient of $(x/r)^6$ simplifies as:

$$\sigma^4(3 - 2\sigma)^2 + 4\sigma^3(1 - \sigma)^3 = \sigma^3[\sigma(3 - 2\sigma)^2 + 4(1 - \sigma)^3].$$

Expanding the bracket:

$$\begin{aligned}\sigma(3-2\sigma)^2+4(1-\sigma)^3 &= \sigma(9-12\sigma+4\sigma^2)+4(1-3\sigma+3\sigma^2-\sigma^3)\\ &= 9\sigma-12\sigma^2+4\sigma^3+4-12\sigma+12\sigma^2-4\sigma^3 = 4-3\sigma.\end{aligned}$$

Therefore the coefficient equals $\sigma^3(4-3\sigma)$, giving:

$$\xi = \varepsilon+\sigma^2(3-2\sigma)\left(\frac{x}{r}\right)^3+\sqrt{\varepsilon^2+2\sigma^2(3-2\sigma)\left(\frac{x}{r}\right)^3\varepsilon+\sigma^3(4-3\sigma)\left(\frac{x}{r}\right)^6}.$$

**Solve for $u$ and $v$**

$$u=\sqrt[3]{4\xi}=2^{2/3}\xi^{1/3}.$$

Using $uv=-p/3=-4(1-\sigma)\sigma(x/r)^2$:

$$v=\frac{uv}{u}=\frac{-4(1-\sigma)\sigma\left(\frac{x}{r}\right)^2}{2^{2/3}\xi^{1/3}}=-2^{4/3}(1-\sigma)\sigma\left(\frac{x}{r}\right)^2\xi^{-1/3}.$$

**Final result**

$$\frac{h_2}{r}=u+v=2^{4/3}\left(\frac{x}{r}\right)^2\sigma(\sigma-1)\,\xi^{-1/3}+2^{2/3}\xi^{1/3},$$

where $\xi$ is given by the expression derived above.

**Equation (9): Phagocytic Capacity $\boldsymbol{k}$**

**Step 1: Simplify $\boldsymbol{a^2 + h_1^2}$**

Using $a^2 = 4\sigma(1-\sigma)x^2$ and $h_1 = 2\sigma x$:

$$a^2 + h_1^2 = 4\sigma(1-\sigma)x^2 + 4\sigma^2 x^2 = 4\sigma x^2[(1-\sigma)+\sigma] = 4\sigma x^2.$$

**Step 2: Expand $\boldsymbol{k}$**

From Eq. (5):

$$4r^2 k \quad = (a^2 + h_1^2) + (a^2 + h_2^2) = 4\sigma x^2 + 4\sigma(1-\sigma)x^2 + h_2^2.$$

Combining the first two terms: $4\sigma x^2 + 4\sigma(1-\sigma)x^2 = 4\sigma x^2(2-\sigma)$. Therefore:

$$k = \sigma(2-\sigma)\left(\frac{x}{r}\right)^2 + \frac{1}{4}\left(\frac{h_2}{r}\right)^2.$$

**Step 3: Substitute $\boldsymbol{h_2/r}$ from Eq. (8) and simplify**

Let $u = 2^{2/3}\xi^{1/3}$ and $v = 2^{4/3}(x/r)^2\sigma(\sigma-1)\xi^{-1/3}$ (so $h_2/r = u + v$). Then:

$$\begin{aligned}
\left(\frac{h_2}{r}\right)^2 \quad &= u^2 + 2uv + v^2 \\
&= 2^{4/3}\xi^{2/3} + 2\cdot 2^{2/3}\xi^{1/3}\cdot 2^{4/3}\left(\frac{x}{r}\right)^2\sigma(\sigma-1)\xi^{-1/3} + 2^{8/3}\left(\frac{x}{r}\right)^4\sigma^2(\sigma-1)^2\xi^{-2/3} \\
&= 2^{4/3}\xi^{2/3} + 2^{(2/3+4/3+1)}\left(\frac{x}{r}\right)^2\sigma(\sigma-1) + 2^{8/3}\left(\frac{x}{r}\right)^4\sigma^2(\sigma-1)^2\xi^{-2/3} \\
&= 2^{4/3}\xi^{2/3} + 8\left(\frac{x}{r}\right)^2\sigma(\sigma-1) + 2^{8/3}\left(\frac{x}{r}\right)^4\sigma^2(\sigma-1)^2\xi^{-2/3}.
\end{aligned}$$

Therefore:

$$\begin{aligned}
k \quad &= \sigma(2-\sigma)\left(\frac{x}{r}\right)^2 + 1/4\left[2^{4/3}\xi^{2/3} + 8\left(\frac{x}{r}\right)^2\sigma(\sigma-1) + 2^{8/3}\left(\frac{x}{r}\right)^4\sigma^2(\sigma-1)^2\xi^{-2/3}\right] \\
&= \sigma(2-\sigma)\left(\frac{x}{r}\right)^2 + 2\left(\frac{x}{r}\right)^2\sigma(\sigma-1) + 2^{-2/3}\xi^{2/3} + 2^{2/3}\left(\frac{x}{r}\right)^4\sigma^2(\sigma-1)^2\xi^{-2/3}.
\end{aligned}$$

Combining the linear terms in $(x/r)^2$:

$$\sigma(2-\sigma) + 2\sigma(\sigma-1) \quad = 2\sigma - \sigma^2 + 2\sigma^2 - 2\sigma = \sigma^2.$$

Hence:

$$k = \sigma^2\left(\frac{x}{r}\right)^2 + 2^{2/3}\left(\frac{x}{r}\right)^4\sigma^2(\sigma-1)^2\,\xi^{-2/3} + 2^{-2/3}\xi^{2/3},$$

where $\xi$ is defined in Eq. (8).

**Equation (10): Normalized Axial Separation $h/r$**

From the definition $h = h_2 - h_1$ and $h_1 = 2\sigma x$:

$$\frac{h}{r} = \frac{h_2}{r} - \frac{h_1}{r} = \frac{h_2}{r} - 2\sigma\left(\frac{x}{r}\right).$$

Substituting Eq. (8) for $h_2/r$:

$$\frac{h}{r} = 2^{2/3}\xi^{1/3} + 2^{4/3}(\sigma - 1)\sigma\left(\frac{x}{r}\right)^2 \xi^{-1/3} - 2\left(\frac{x}{r}\right)\sigma,$$

where $\xi$ is defined in Eq. (8).

**Equation (11): Cubic Relation among $h/r$, $k$, and $\varepsilon$**

This key result is derived by eliminating $h_1$, $h_2$, and $a$ from Eqs. (4) and (5), retaining only $h = h_2 - h_1$, $k$, $\varepsilon$, and $r$.

**Step 1: Express $a^2$ in terms of $k$, $h_1$, $h_2$**

From Eq. (5):

$$4r^2k = (a^2 + h_1^2) + (a^2 + h_2^2) = 2a^2 + h_1^2 + h_2^2.$$

Solving for $a^2$:

$$2a^2 = 4r^2k - h_1^2 - h_2^2.$$

**Step 2: Expand the volume expression**

From Eq. (4), multiply through by $8r^3$:

$$8r^3\varepsilon = h_2(3a^2 + h_2^2) - h_1(3a^2 + h_1^2) = 3a^2(h_2 - h_1) + (h_2^3 - h_1^3).$$

Using $h_2^3 - h_1^3 = (h_2 - h_1)(h_2^2 + h_1h_2 + h_1^2)$ and $h = h_2 - h_1$:

$$8r^3\varepsilon = 3a^2\,h + h\,(h_2^2 + h_1h_2 + h_1^2).$$

**Step 3: Substitute the expression for $a^2$**

$$\begin{aligned}
8r^3\varepsilon &= 3 \cdot \frac{4r^2k - h_1^2 - h_2^2}{2} \cdot h + h\,(h_2^2 + h_1h_2 + h_1^2) \\
&= h\left[\frac{3(4r^2k - h_1^2 - h_2^2)}{2} + h_2^2 + h_1h_2 + h_1^2\right] \\
&= h\left[6r^2k - \frac{3}{2}h_1^2 - \frac{3}{2}h_2^2 + h_1^2 + h_1h_2 + h_2^2\right] \\
&= h\left[6r^2k - \frac{1}{2}(h_1^2 - 2h_1h_2 + h_2^2)\right] \\
&= h\left[6r^2k - \frac{1}{2}(h_2 - h_1)^2\right] \\
&= h\left[6r^2k - \frac{h^2}{2}\right].
\end{aligned}$$

**Step 4: Non-dimensionalize and rearrange**

Dividing by $r^3$:

$$8\varepsilon = \frac{h}{r}\left[6k - \frac{1}{2}\left(\frac{h}{r}\right)^2\right] = 6k\left(\frac{h}{r}\right) - \frac{1}{2}\left(\frac{h}{r}\right)^3.$$

Multiplying by $-2$:

$$\left(\frac{h}{r}\right)^3 - 12k\,\frac{h}{r} + 16\varepsilon = 0.$$

This is a depressed cubic in $h/r$ with coefficients $p = -12k$ and $q = 16\varepsilon$. Crucially, *this equation does not contain $x$ or $\sigma$*: the normalized axial separation depends only on $k$ and $\varepsilon$, independently of target size and curvature.

**Discriminant and solution structure**

The discriminant is:

$$\delta = -4p^3 - 27q^2 = -4(-12k)^3 - 27(16\varepsilon)^2 = 6912k^3 - 6912\varepsilon^2 = 6912(k^3 - \varepsilon^2).$$

The physical constraint $\varepsilon^2 \leq k^3$ (cell surface cannot decrease below the sphere equivalent of the remaining volume) ensures $\delta \geq 0$, so Eq. (11) admits three distinct real roots: two positive and one negative (inadmissible) for frustrated phagocytosis geometries.

### Generalization of Eq. (11) to a Flat Target

The manuscript states (Figure 5E) that the same cubic relation holds when the phagocytic target is a flat surface. Here we derive this result from first principles.

#### Geometry

When the target is flat, the phagosome is a flat disk of radius $a$, and the free cell surface is a spherical cap of height $h$ and base radius $a$ sitting above the flat substrate. The quantity $h$ is now the axial distance from the flat substrate to the apex of the free cell surface, identical to the definition in Figure 5E of the main text.

#### Surface area and volume

The total post-phagocytosis surface area consists of:

- Phagosome (flat disk): $\pi a^2$
- Free surface (spherical cap of height $h$): $\pi(a^2 + h^2)$

so the phagocytic capacity is:

$$k = \frac{\pi a^2 + \pi(a^2 + h^2)}{4\pi r^2} = \frac{2a^2 + h^2}{4r^2}.$$

The post-phagocytosis cell volume is that of the spherical cap alone:

$$\varepsilon = \frac{1/6\,\pi h(3a^2 + h^2)}{4/3\,\pi r^3} = \frac{h(3a^2 + h^2)}{8r^3}.$$

#### Elimination of $a^2$

From the expression for $k$:

$$2a^2 = 4kr^2 - h^2 \quad \Rightarrow \quad 3a^2 = 6kr^2 - 3/2\,h^2.$$

Substituting into the volume equation:

$$8\varepsilon r^3 \quad = h(3a^2 + h^2) = h(6kr^2 - 3/2\,h^2 + h^2) = h(6kr^2 - 1/2\,h^2).$$

Multiplying by 2:

$$16\varepsilon r^3 = 12kr^2 h - h^3.$$

Dividing by $r^3$:

$$\left(\frac{h}{r}\right)^3 - 12k\left(\frac{h}{r}\right) + 16\varepsilon = 0,$$

which is identical to Eq. (11). This confirms that the geometric criterion for nuclear involvement is the same for a flat target as for a convex spherical target, independently of target curvature.

**Generalization of Eq. (11) to a Concave Spherical Target**

The manuscript (Figure 5F) also asserts that Eq. (11) holds for a concave spherical substrate. Below we derive this case, which features a sign change in the volume expression.

**Geometry**

When the target is a concave spherical surface (e.g., the inner surface of a sphere of radius $x$), the phagosome cap of height $h_1$ opens *inward*, toward the center of the concavity. In this configuration, both the phagosome cap and the free-surface cap lie on the *same* side of the contact plane, so they each *add* positively to the cell volume. The axial separation $h$ is defined as $h = h_1 + h_2$ (the sum, not the difference), representing the total axial extent of the cell between the two cap apices.

**Modified surface area and volume definitions**

The phagocytic capacity retains the same form as before:

$$k = \frac{\pi(a^2 + h_1^2) + \pi(a^2 + h_2^2)}{4\pi r^2} \quad \Rightarrow \quad 2a^2 + h_1^2 + h_2^2 = 4kr^2 \qquad \text{(C1)}$$

The volume now sums both caps:

$$\varepsilon = \frac{1/6\,\pi h_2(3a^2 + h_2^2) + 1/6\,\pi h_1(3a^2 + h_1^2)}{4/3\,\pi r^3} \quad \Rightarrow \quad h_2(3a^2 + h_2^2) + h_1(3a^2 + h_1^2) = 8\varepsilon r^3 \qquad \text{(C2)}$$

**Derivation**

**Step 1.** Expand $h = h_1 + h_2$ to get $h^2 = h_1^2 + h_2^2 + 2h_1h_2$. Combining with (C1):

$$h_1^2 + h_2^2 = 4kr^2 - 2a^2, \qquad 2h_1h_2 = h^2 - (h_1^2 + h_2^2) = h^2 - 4kr^2 + 2a^2 \qquad \text{(C3)}$$

**Step 2.** Factor (C2):

$$(h_1 + h_2) \cdot 3a^2 + h_1^3 + h_2^3 = 8\varepsilon r^3.$$

Use $h_1^3 + h_2^3 = (h_1 + h_2)(h_1^2 - h_1h_2 + h_2^2)$:

$$h[3a^2 + h_1^2 + h_2^2 - h_1h_2] = 8\varepsilon r^3.$$

**Step 3.** Substitute using (C1) and (C3):

$$\begin{aligned} h_1^2 + h_2^2 &= 4kr^2 - 2a^2, \\ h_1h_2 &= 1/2\,(2a^2 + h^2 - 4kr^2) = a^2 + h^2/2 - 2kr^2. \end{aligned}$$

Therefore:

$$h_1^2 + h_2^2 - h_1 h_2 \quad = (4kr^2 - 2a^2) - (a^2 + h^2/2 - 2kr^2) = 6kr^2 - 3a^2 - h^2/2\,.$$

**Step 4.** Insert into the factored volume equation:

$$\begin{aligned} h[3a^2 + 6kr^2 - 3a^2 - h^2/2] &= 8\varepsilon r^3 \\ h[6kr^2 - h^2/2] &= 8\varepsilon r^3 \\ 12kr^2h - h^3 &= 16\varepsilon r^3. \end{aligned}$$

Dividing by $r^3$:

$$\left(\frac{h}{r}\right)^3 - 12k\left(\frac{h}{r}\right) + 16\varepsilon = 0,$$

which is again identical to Eq. (11). This result confirms that the geometric criterion for nuclear deformation during frustrated phagocytosis is invariant to the sign (polarity) of the substrate curvature.

**Equation (12): Physically Relevant Root of Eq. (11)**

**Three real roots via the trigonometric method**

When $\delta > 0$ (i.e., $k^3 > \varepsilon^2$), the three real roots of $t^3 - 12k\,t + 16\varepsilon = 0$ are given by the trigonometric formula. With $p = -12k$:

$$t_j = 2\sqrt{\frac{-p}{3}}\cos\left(\frac{1}{3}cos^{-1}\left(\frac{3q}{2p}\sqrt{\frac{-3}{p}}\right) - \frac{2\pi j}{3}\right), \quad j = 0{,}1{,}2.$$

Substituting $p = -12k$ and $q = 16\varepsilon$:

$$\begin{aligned} \sqrt{-p/3} &= \sqrt{4k} = 2\sqrt{k}, \\ \frac{3q}{2p}\sqrt{\frac{-3}{p}} &= \frac{3 \cdot 16\varepsilon}{2(-12k)} \cdot \frac{1}{2\sqrt{k}} = \frac{-\varepsilon}{k^{3/2}}. \end{aligned}$$

Using $cos^{-1}\left(-\varepsilon/k^{3/2}\right) = \pi - tan^{-1}\left(\sqrt{k^3/\varepsilon^2 - 1}\right)$, the three roots are:

$$t_j = 4\sqrt{k}\cos\left(\frac{\pi}{3} - \frac{1}{3}tan^{-1}\left(\sqrt{\frac{k^3}{\varepsilon^2} - 1}\right) - \frac{2\pi j}{3}\right).$$

**Identifying the physically relevant root**

Of the three roots, $j = 2$ is always negative and therefore physically inadmissible. The remaining two roots, $j = 0$ and $j = 1$, are always positive but respond oppositely to $k$: the $j = 0$ root increases monotonically with $k$, while the $j = 1$ root decreases monotonically with $k$. Because a larger membrane capacity permits deeper engulfment and thereby reduces the axial clearance, the physically relevant solution must have $h/r$ decreasing with increasing $k$. This condition uniquely identifies the $j = 1$ root:

$$\begin{aligned} \frac{h}{r} &= 4\sqrt{k}\cos\left(\frac{\pi}{3} - \frac{1}{3}tan^{-1}\left(\sqrt{\frac{k^3}{\varepsilon^2} - 1}\right) - \frac{2\pi}{3}\right) \\ &= 4\sqrt{k}\cos\left(-\frac{\pi}{3} - \frac{1}{3}tan^{-1}\left(\sqrt{\frac{k^3}{\varepsilon^2} - 1}\right)\right) \\ &= 4\sqrt{k}\cos\left(\frac{\pi}{3} + \frac{1}{3}tan^{-1}\left(\sqrt{\frac{k^3}{\varepsilon^2} - 1}\right)\right), \end{aligned}$$

where the last step uses $\cos(-\theta) = \cos(\theta)$ and the identity $\cos(\pi/3 + \theta) = \cos(-\pi/3 - \theta)$. Therefore:

$$\frac{h}{r} = 4\sqrt{k}\cos\left(\frac{\pi}{3} + \frac{1}{3}\tan^{-1}\sqrt{\frac{k^3}{\varepsilon^2} - 1}\right).$$

**Consistency with flat and concave targets**

Equation (11) contains no target radius $x$, confirming that Eq. (12) holds identically when the phagocytic target is a flat surface (formally $x \to \infty$) or a concave spherical surface, as demonstrated in Figure 5E–F of the main text. The onset of nuclear deformation is therefore independent of target curvature.

**Equation (13): Phagocytic Capacity for Multiple Small Targets**

When a phagocyte engulfs $n$ identical small spherical targets of radius $x$, the post-phagocytosis cell consists of the sphere-equivalent body from which $n$ spherical caps have been replaced by phagosome caps, each of area $4\pi x^2$ and volume $4/3\,\pi x^3$.

**Surface area ratio**

The post-phagocytosis surface area is the free outer surface area plus $n$ phagosome areas. Assuming a surface-minimizing cell geometry and volume ratio $\varepsilon$, the outer surface becomes a sphere of adjusted radius $r'$ where $(r')^3 = r^3\varepsilon + nx^3$. The total surface area is:

$$S_{\mathrm{post}} = 4\pi(r')^2 + 4\pi n x^2 = 4\pi\left(\varepsilon + n\left(\frac{x}{r}\right)^3\right)^{2/3} r^2 + 4\pi n x^2.$$

Dividing by $4\pi r^2$:

$$k = \left(\varepsilon + n\left(\frac{x}{r}\right)^3\right)^{2/3} + n\left(\frac{x}{r}\right)^2.$$

This expression (derived in Ref. [14] of the main text) enables determination of the intrinsic phagocytic capacity $k_{\mathrm{intrinsic}}$ from experiments in which macrophages are allowed to engulf many small particles until saturation, thereby revealing the membrane-limited capacity free of nuclear constraints.

**Summary Table of Derived Equations**

| Eq. | Quantity | Key derivation step |
|---|---|---|
| (1) | $a^2 + (h_1 - x)^2 = x^2$ | Pythagorean theorem, target sphere |
| (2) | $a^2 + (h_2 - R)^2 = R^2$ | Pythagorean theorem, free-surface sphere |
| (3) | $\sigma = h_1/(2x)$ | Spherical cap area fraction |
| (4) | $\varepsilon$ | Difference of spherical cap volumes |
| (5) | $k$ | Sum of spherical cap surface areas |
| (6) | $h = h_2 - h_1$ | Definition |
| (7) | Cubic in $h_2/r$ | Substitution of (1),(3) into (4) |
| (8) | $h_2/r$ (closed form) | Cardano's formula applied to (7) |
| (9) | $k$ (closed form) | Substitution of (8) into (5) |
| (10) | $h/r$ (closed form) | Combining (8) and (6) |
| (11) | Cubic in $h/r$, $k$, $\varepsilon$ | Eliminating $h_1, h_2, a$ from (4),(5) |
| – | Same cubic, flat target | Flat phagosome; $h =$ cap height |
| – | Same cubic, concave target | Volume sum; $h = h_1 + h_2$ |
| (12) | $h/r$ from $k$, $\varepsilon$ | Trigonometric root of (11) |
| (13) | $k$ for $n$ small targets | Volume/area accounting for $n$ phagosomes |
